# Microplate micromilling: A customizable platform to support the prototyping, development and testing of microphysiological culture models

**DOI:** 10.1101/2024.10.02.615399

**Authors:** Jacob I. Reynolds, Meredith Adams, Jose Jimenez, Brian P. Johnson

**Author notes:** denotes corresponding author. Electronic Supplementary Information (ESI) available.

## Abstract

The development of microphysiological cell culture models (MPMs) that align with the throughput demands of drug and chemical testing are needed to help reduce animal testing, aide in the discovery of new drugs, and identify harmful chemical exposures. To address this need, we have developed a process for rapid prototyping MPM devices using computer numerical control (CNC) micromilling of commercially available microplates. Microchannels are cut out of the existing microplate structure and ports are drilled into the bottom of the wells to interface the wells. To test versatility and benchmark to another rapid-prototyping approach, we manufactured common microfluidic features into microplates using four different CNC mills as well as a 3D printer. Cell viability was assessed for polystyrene (PS) well plates and two 3D printed resins (MED610 and VeroClear) with the PS showing >2.5-fold increase in cell growth after three days. Machines were tested on their ability to create common device features including a traditional microfluidic device as well as a custom design incorporating complex geometries. Features were measured by confocal microscopy. We found that features including 1000µm ports, 800µm microchannels, 200µm phase-guides, and 500µm post arrays were machined and the range of CVs for features were 1.02-4.42, 1.32-3.50, 2.34-16.58, 6.25-16.40 respectively, while the 3D printed features exhibited maximal CVs of 20.98, 11.68, 23.60, and 10.01 for the same features. Predictably, more expensive machines generally showed higher accuracy and lower variation, but many features could be created accurately and precisely by inexpensive (<$3000) machines facilitating the broader use of this technology to create a user customizable platform to support the prototyping, development, and testing of human relevant models with broad applications across the life sciences.

**Graphical abstract:** Multiple CNC mills are assessed on accuracy and precision of microfluidic features of interest for microphysiological model development creation.

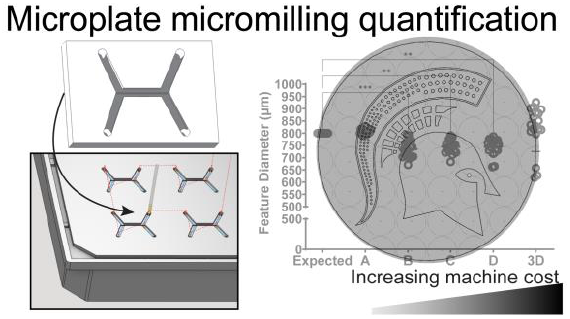

## Introduction

The properties of fluids at the microscale are useful in creating microphysiological models (MPMs) and enable the creation of cell culture models that better mimic natural physiology as compared to traditional 2D cell culture. The use of small volumes allows for low reagent consumption while also reducing the use of limited test compounds or hazardous agents ^1-3^. Microfluidic devices have a high surface area to volume ratio that facilitates fast mass and heat transfer^1, 4^-^5, 6^.There have been numerous advancements in advanced cell culture technologies, however these advanced technologies also come with a set of challenges that make their application for drug and chemical screening applications problematic. These challenges include complicated pump/valve systems, low throughput capacity, and incompatible materials, namely polydimethylsiloxane (PDMS). Challenges aside, these technologies are being applied to a wide range of fields leading to important scientific discoveries. To support drug discovery and chemical screening during the shift away from animal models there is a need for more biologically relevant cell culture models that maintain the throughput compatibility of traditional microplate-based formats. The manufacture of plate-based devices from biocompatible materials is one way to help address these challenges and promote ease of adoption.

Rapid prototyping of cell culture devices is possible via computer numerical control (CNC) micromilling or 3D printing, but CNC machining holds several advantages ^6^ (reviewed in^7^). Microplate micromilling uses subtractive manufacturing to create microscale features (<1000µm) either into the top or bottom of a standard polystyrene (PS) microplate as depicted in **Figure 1**^8-10^. A unique novelty of the microplate micromilling technology is that unlike many microfluidic/microphysiological systems, these devices are manufactured from commercially available injection molded American National Standards institute (ANSI) microplates, providing a familiar platform for new users and eliminating fabrication time and cost for the base plate. Fluidics can be incorporated into multiple plate formats including 6, 24, 48, 96, and 384 well formats. Since the plate format is preserved, these devices keep their compatibility with common lab equipment including liquid handlers, plate readers, microscopes, shakers, etc. Integration with laboratory equipment along with their PS construction makes these devices well suited for cell culture applications and assay development. The use of PS also overcomes the problem of hydrophobic molecules being sequestered by the device, which is a problem commonly seen in typical microfluidic devices constructed from PDMS ^11, 12^. Eliminating the sequestration of lipophilic molecule s makes these devices well suited for drug and chemical screening.

**Figure 1.**
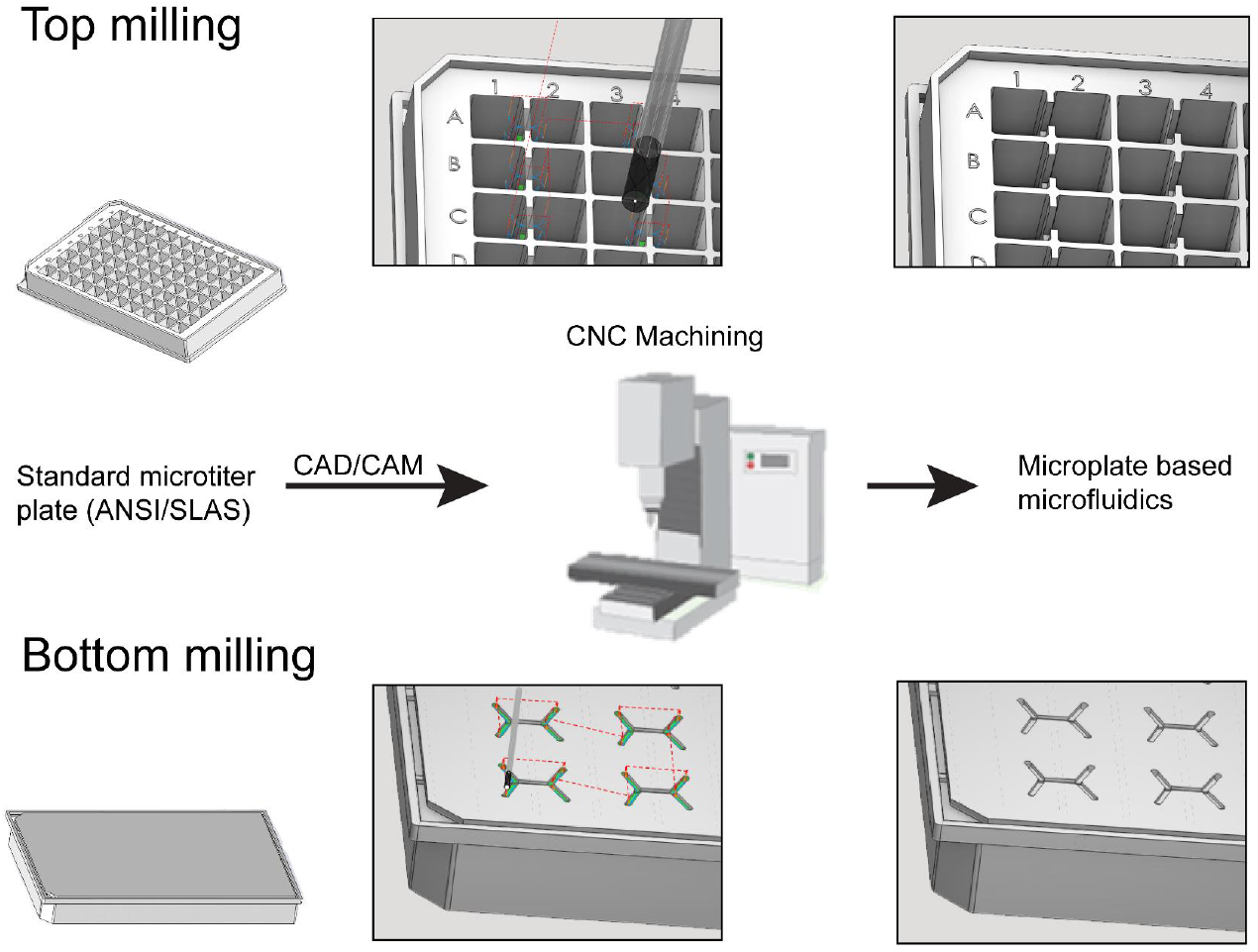
Overview of CNC machining of microtiter plates. CAD and CAM software is used to generate the designs that the CNC machine cuts into the plate. These devices can then be used for cell culture and other microfluidic application while maintaining the usability of the well plate format.

While we have successfully employed CNC micromachining of devices into well plates for specific use cases the broader use as a platform technology to create a customizable platform to support the prototyping, development and testing of human relevant models with broad applications across the life sciences has not fully been explored^8-10, 13^. The adoption of micromilling to the life science community has been hindered by several barriers of adoption. Professional grade CNC machines are expensive and large, requiring both substantial capital, lab space, and expertise to operate. With these large machines comes other concerns including noise level, vibrations, and a factor of intimidation to those unfamiliar with heavy machinery. The increased availability of hobbyist grade CNC addresses these potential hurdles of adoption and presents a potential avenue to labs looking to adopt micromilling. Creating the CAD models and using them to create code to create the machines offers another potential barrier of entry. The ecosystem of CAD and CAM software has evolved to now include simple to learn and free options. Training through included tutorials and online resources can give a motivated individual the skillset to create their own models and machine code. An important consideration for designing devices for microplate micromilling is that only the features that need to be cut need to be modelled. This means that often it is not necessary to model the entire plate and all its’ geometry. For example, when designing a device to be cut into the bottom of the plate, the geometry of the wells is irrelevant save for the relation of spacing for port holes and channel orientation.

In this work, a set of commonly used microscale features of interest in MPM creation were arranged in both a simple and complex geometric pattern and manufactured using four commercially available CNC machines ranging from hobbyist to professional grade, as well as a professional grade resin 3D printer. From this work, it was found that microfluidic features down to 200µm could be accurately manufactured using CNC machines. Accuracy and precision of features was found to correlate with the grade of machine with the professional grade mill accurately producing all features. Even the hobbyist grade mill that performed the worst of the mills tested produced 1000µm ports (927.4±44.3µm), 800µm channels (783.4±29.6µm), and 200µm phase-guides (155.4±20.3µm) arranged in a simple geometric pattern that would likely satisfy tolerance requirements. The precision of the hobbyist mill was similarly impressive with a range of CVs from 2.03-12.12 (1000µm channel and 200µm phase-guide respectively). The accuracy and precision found for all tested machines demonstrates the robustness of micromilling and the feasibility of the technology to be implemented to in the creation of user customizable MPMs with a hobbyist grade machine.

## Experimental

### Methods

Devices and well plates were modelled in CAD software ((Solidworks, Dessault Systems, Vélizy-Villacoublay, France). Tool paths were generated using CAM software and processed to G-code using a MACH3 (mills B, C, D) or Fanuc(30i) (mill A) postprocessor (SprutCAM America, Waunakee, WI, USA) and are included in **Supplemental File 1**. Feeds, speeds, and milling strategy were kept consistent between mills and were selected off of previous work performed by Guckenberg et al ^**6**^. Plates were milled using an equidistant strategy with both climb and conventional, 50% stepover, 50% depth step, feed of 254mm/min, and speed of 5000RPM. Milling was performed using a Hurco VM20 (Mill A), Syil X5 Combo (Mill B), Syil X5 Speedmaster (Mill C), and a Baleigh DEM-0906 (Mill D). Devices were milled into the bottom of clear Corning Falcon 96 (REF 353072 Lot 3015014) polystyrene (PS) well plates. The well plates were secured directly to the deck of the mill using t-channel press down clamps for mills C and D. A vise was used to secure the plate on mills A and B. Test devices were milled using an 800µm 2FL SQ TA endmill (Kyocera SGS 20365) and a 500µm 2FL SQ TA endmill (Kyocera SGS 02362). To set the Z origin a piece of paper (i.e. post it note) was used to determine the relation between the end of the endmill and the surface of the plate. Briefly, the mill was stepped towards the surface of the plate at a fine resolution (25µm) with the paper under the endmill. Moving the paper back and forth allows the operator to feel as the endmill begins to catch and drag on the paper. As soon as the tool comes down enough to fully stop the paper from moving the operator knows that the end of the endmill is one thickness of paper above the surface of the workpiece. Finding this point and accounting for the thickness of the paper allows the operator to correctly set the origin. A correction of 75µm was applied for this study. Six plates were milled for each machine. For mill A a single trial of 6 plates were made. For mills B-D 2 trials of 3 plates each were made. Prior to imaging, milled plates were sonicated in 100% isopropanol (IPA) for 15 minutes, washed with DI water, and air dried. Images for each feature were obtained using a standardized measurement protocol using a confocal imaging system (Yokogawa CQ1). Measurements were made using image analysis using ImageJ software with each feature was measured in three different locations ^14^. All measurements are included in **Supplemental File 2**.

3D printed devices were printed using a Stratasys J750 PolyJet printer through the 3D Printing Core of the Institute of Quantitative Health Science and Engineering at Michigan State University. Microfluidic devices and well plates were modelled in CAD software ((Solidworks, Dessault Systems, Vélizy-Villacoublay, France). Solid models were converted to a mesh and sliced using GrabCAD (Stratasys). Devices were printed using either VeroClear (Stratasys) or MED610 resin (Stratasys). Support material was removed through physical manipulation and use of a water cabinet. Printed plates were sonicated in 100% IPA for 15min, dried with compressed air, and plasma treated with Oxygen gas plasma for 5min, and sterilized using UV prior to cell culture.

The Corning 96 well plates used for the mill testing (REF 353072 Lot 3015014) were assessed for flatness. Plates were imaged on the same confocal imaging system with a 10X objective (Yokogawa CQ1). The relative z position obtained using the autofocus (AF) wave measurement and recording the motor position for the peak. These values were averaged and the difference between them taken to be the average depth of the milling. The AF wave measurement tool was used to collect the relative z position for the five locations across the plate (wells A1, H1, H12, A12, E6) and at seven locations expanding across a single well (E7). The difference in flatness across the plate and within a well was found by taking the difference of the max and min values of the five or seven data points respectively. Three plates were assessed for flatness.

HepG2 cells were cultured and maintained using standard protocols (ATCC). Cells were dispensed using a Biotek liquid handler (Agilent). Cell seeding density was adjusted for surface area and seeded at 10,000 per well for the printed plates and 5,000 per well in the commercial plate (Corning Falcon 96 (REF 353072 Lot 3015014). Cells were grown in a standard humidified incubator under 5% CO2. Viability was assessed at 24, 48, and 72h using CellTiter-Glo 2.0 (Promega) as directed. Briefly, CellTiter-Glo 2.0 reagent as added in equal volume to the media in the well after removing half the media. Plates were shaken at 300 RPM for 2 minutes then incubated at room temperature for 10 minutes to stabilize signal prior to reading on a plate reader. Signal was normalized to the area of the well. A second study was conducted using identical culture conditions to assess imaging ability of the plates. On day 4, cells were stained with Hoechst 1:1000, Mitotracker red 1:500, and Calcein AM 1:1000 in media and incubated for 20 minutes. Cells were imaged using a confocal imaging system (Yokogawa CQ1) using a 20X objective. Images are displayed using a standardized lookup table and image histograms for each of the four channels can be found in **Supplemental Figure 1**.

Noise measurements were collected using a calibrated Cirrus Sound Level Meter, Type2, set for “A” weighting, slow response. The sound level meter was calibrated according to the manufacturer’s specifications. Locations were selected based on concerns and to be as representative as possible. Testing was performed a CNC machine (Tormach 1100) in both the operating and off conditions. A map of the noise recordings is provided in **Supplemental File 3**.

To assess for differences between trials, two-way ANOVA followed by Sidak’s multiple comparisons test was performed using GraphPad Prism version 10.1.2 (GraphPad software). Measurements for each feature from each of the six plates per mill were pooled and used for analysis between machines. To assess the accuracy of each machine, one-way ANOVA followed by Tukey’s multiple comparison test was performed using GraphPad Prism version 10.1.2 (GraphPad Software). The expected value is defined to be the nominal value assigned to the feature during the CAD modelling (e.g., 800µm for the 800µm wide channels). To assess the precision of each machine, row statistics for mean and coefficient of variation (%CV) were performed using GraphPad Prism version 10.1.2 (GraphPad Software). Significance is reported as *P⩽0.05, **P⩽0.01, ***P⩽0.001, ****P⩽0.0001.

## Results & discussion

Material biocompatibility is a major consideration when creating MPMs, so we first set out to validate material biocompatibility. The testing of the 3D printing resins demonstrate that special care should be taken when using these resins to create cell culture devices. For the 3-day viability study, the standard PS plate was found to have increased Cellglo luminescence corresponding to a higher cell number for all time points as compared to Veroclear and MED610 as shown in **Figure 2A**. Cell growth was found in the cells of the PS plate as noted by increasing cellular ATP as detected by Cellglo signal over time. Cellular ATP levels in cells grown in both 3D resins remained steady over time suggesting that cells remain alive but are not expanding as they are in the PS plates due to some influence of the resin. The ability to image cells is also a major consideration for New Approach Methodologies (NAMs) development. Challenges imaging through both the Veroclear and MED610 were noted during testing for both brightfield and fluorescent microscopy (**Figure 2C-D**). In brightfield, the presence of printing remnants and potential variations in flatness made focusing on the cells problematic. The fluorescent staining tested exhibited low signal in both the Veroclear (**Figure 2D**) or MED610 (**Figure 2C**). We hypothesize that dyes may be interacting with/partitioning to the resins and are not making it to the cells at a high enough concentration. Using the same dyes and staining protocol produced clear images for the PS plate (**Figure 2B**).

**Figure 2:**
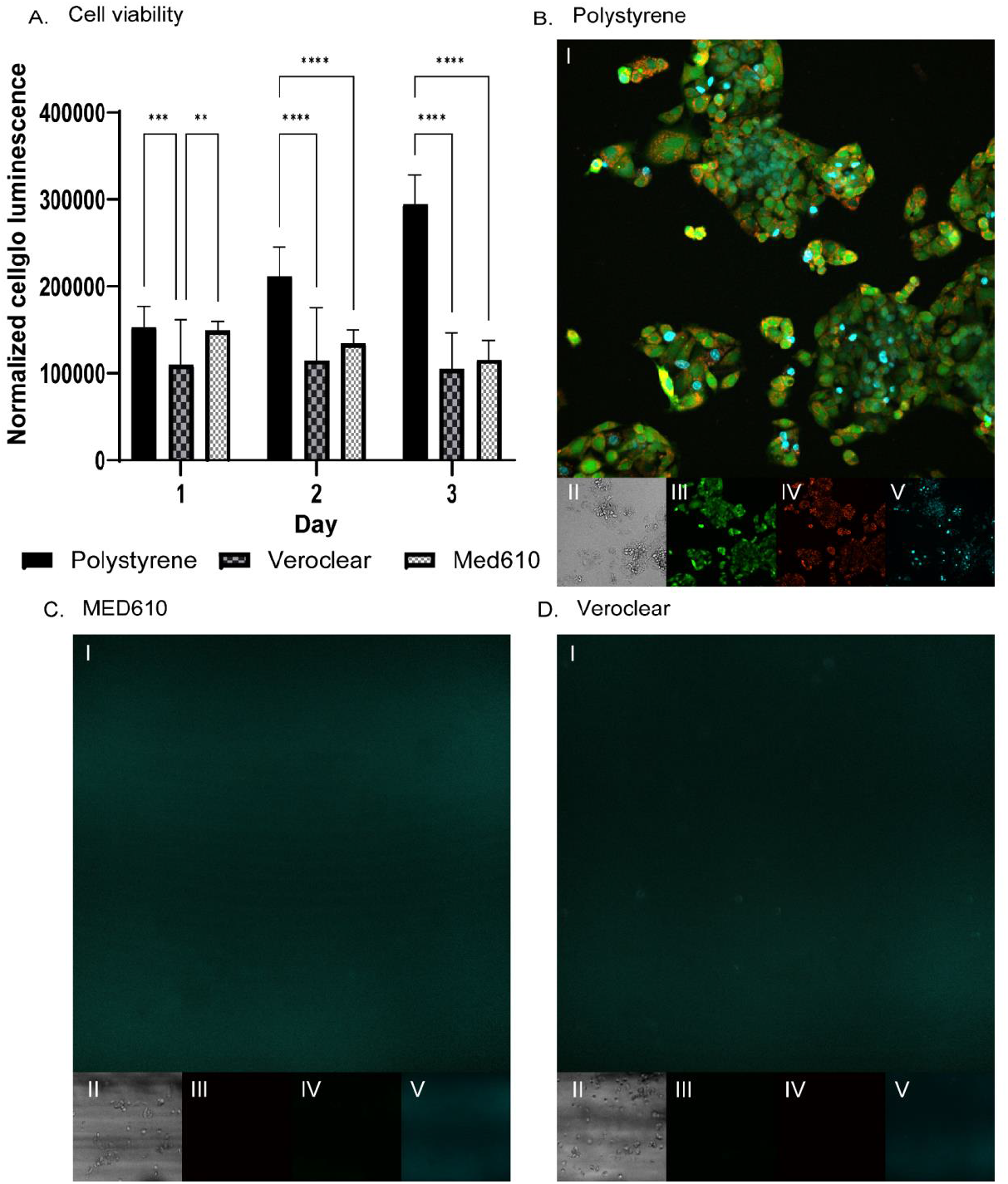
**A)** Plot of cell viability using CellTiter Glo luminescence. Significance is reported as *P⩽0.05, **P⩽0.01, ***P⩽0.001. **B-D)** 20X microscopy images of day 4 HepG2 cells **B)** Polystyrene, **C)** MED610, **D)** Veroclear. **I)** fluorescent composite, **II)** brightfield, **III)** cell body, **IV)** mitochondria, **V)** nucleus

Bottom milling incorporates microfluidic devices into the bottom of the microplate. This allows both existing and novel designs to be incorporated into a well plate format. For example, a traditional PDMS phase separation device is depicted in **Figure 3A**. To evaluate the ability of micromilling to recreate traditional microfluidic devices with simple geometry a traditional phase separation device was designed into the bottom of a 96 well plate as depicted in **Figure 3B**. This device was designed to test the capabilities of the machines to create channels and ports without complex geometry or curves.

**Figure 3:**
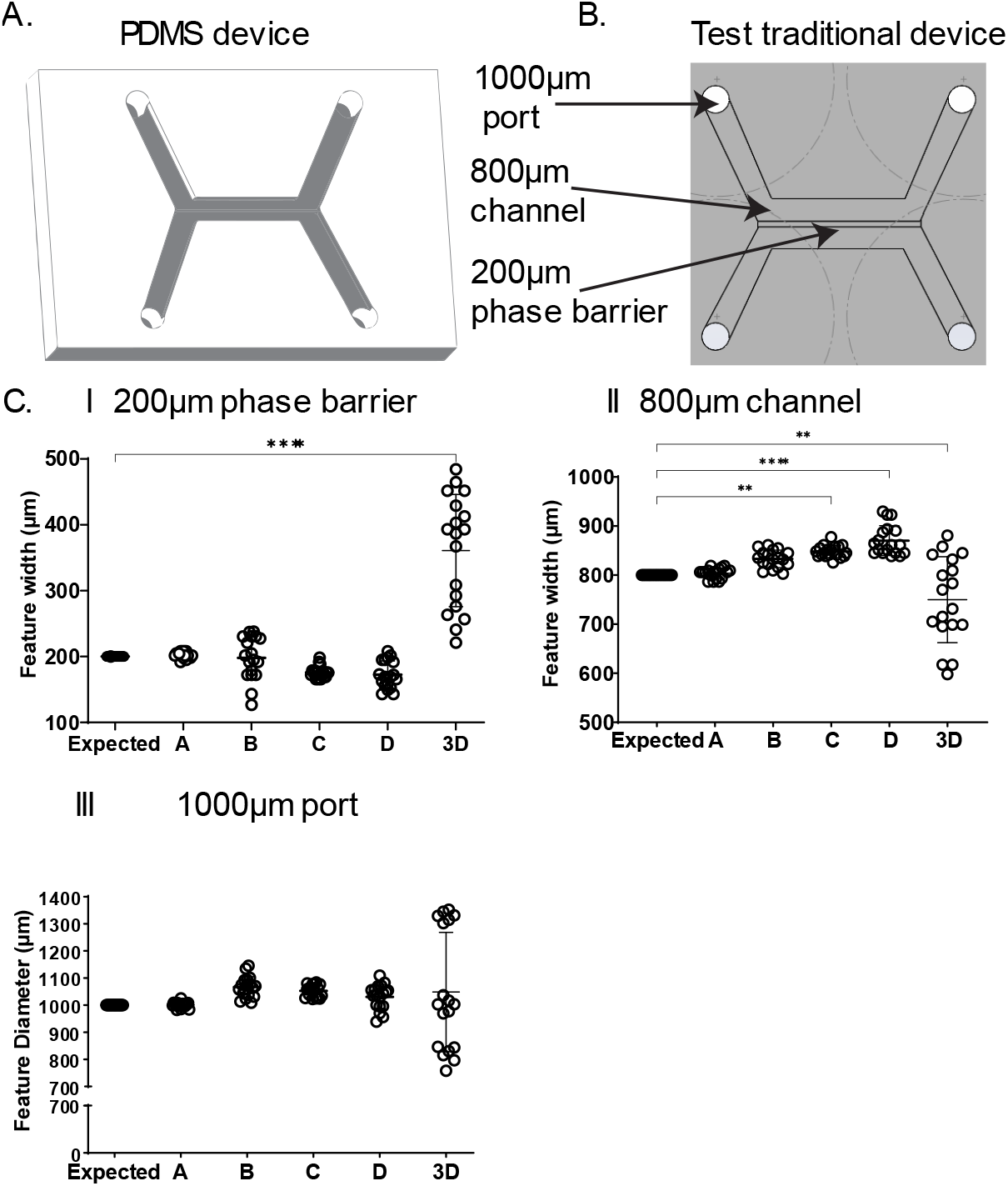
**A)** Model of traditional Y type microfluidic device. **B)** Detailed view of Y device with features of interest labelled. **C)** Measurements of milled microfluidic features for the traditional device. Significance is reported as *P⩽0.05, **P⩽0.01, ***P⩽0.001.**I)** 200µm phase barrier **II)** 800µm channel **III)** 1000µm port

The results of the traditional device test with the four tested mills demonstrate that CNC micromilling is a viable manufacturing technique to create traditional microfluidic devices directly into standard well plates. Six technical replicates (ie plates) with three measurements per feature (n=18) are reported for each machine. The data for the traditional device for all machines tested appear in **Figure 3C**. The 200µm phase barrier, was cut accurately for mills A, B,C, D, but not by the 3D printer (means of 202.8µm, 178.4µm, 158.4µm, 155.4µm, 361.0µm (P <0.0001) respectively. The 800µm channel was cut accurately for mills A and B with significant differences found for C, D, and 3D (means of 803.7µm, 833.1µm, 848.3µm (P=0.0046), 870.3µm (P <0.0001), 749.8µm (P=0.0028), respectively. The 1000µm diameter port was cut accurately for all machines tested (means of 1000.6µm, 1066.8µm, 1052.8µm, 1030.3µm, and 1048.3µm). The results of the traditional test demonstrate the ability of CNC machining to accurately create simple microfluidic features directly into a well plate. The tolerances that can be achieved vary with machine and correlate with grade of machine. It should be noted that even the features that were found to be statistically different from expected may not pose an issue to a potential adoptee. For example, if a user had a device with a 800µm channel with a bilateral 50µm tolerance, mills A, B, and C would satisfy this requirement even though C was found to be statistically different. The numbers and statistics presented here are intended to serve as a starting point of what can be expected from these machines and not what is or is not acceptable in device manufacture.

To test the capabilities of the machines more thoroughly, a complex geometry device comprised of pillar arrays and channels of varying dimensions arranged in complex curves was designed. **Figure 4A** details the features of interest for the device. The measurements and standard deviations for the features in the complex geometry device as well as any significant differences from expected are reported in **Figure 4B**. The 1000µm pillars were made accurately except for mills B and C (means of 997.4µm, 942.6µm (P=0.0001), 949.8µm (P<0.0001), 974.5µm, 1030.6µm respectively). For the 800µm pillar mill A and the 3D printed created accurate features while mills B, C, and D were significantly different from expected (means of 806.9 µm, 734.5µm (P=0.0004), 744.6µm (P=0.0042), 744.6µm (P=0.0042), 818.2µm respectively). All the machines tested except for mill A measured significantly different from expected for the 500µm pillars (means of 471.2µm, 380.5µm (P<0.0001), 431.9µm (P=0.0001), 454.0µm (P=0.0247), 594.7µm (P<0.0001). The 1000µm channel was made accurately except for mill D (means of 1000.3µm, 1083.6µm, 1045.0µm, 1143.9µm (P=0.0037), 986.9µm). The same trend was found for the 800µm channels (means of 819.9µm, 796.5µm, 776.5µm, 837.0µm (P=0.0305), 723.2µm respectively). The 500µm channel was cut accurately by mill A with all other machines measuring significantly different from expected (means of 539.0µm, 611.0µm (P<0.0001), 554.5µm (P=0.0069), 602.2µm (P<0.0001), 346.4µm (P<0.0001) respectively). The differences between individual machines are of potential interest to potential adoptees of microplate micromilling. In **Supplemental Table 2** all the pairwise comparisons between the 5 machines tested in this study for both the traditional and complex devices are included.

**Figure 4:**
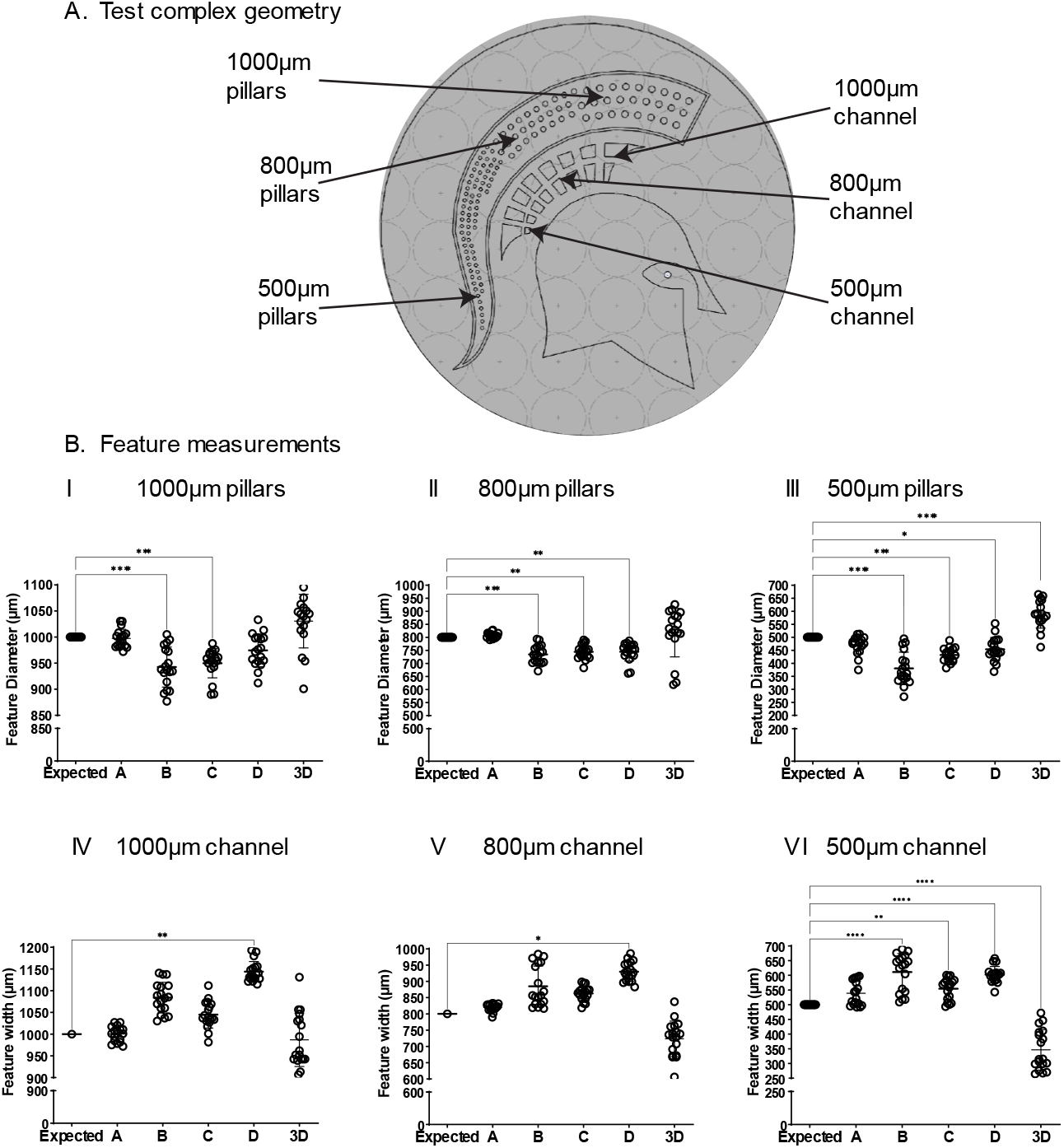
**A)** Detailed view of complex geometry device with features of interest labelled. **B)** Measurements of features for the complex geometry device. Significance is reported as *P⩽0.05, **P⩽0.01, ***P⩽0.001. **I)** 1000µm pillar **II)** 800µm pillar **III)** 500µm pillar **IV)** 1000µm channel **V)** 800µm channel **VI)** 500µm channel **VII)** coefficient of variation

The precision of the tested machines for the features of the traditional and complex devices was calculated as the coefficient of variation (CV) is reported in **Table 1**. Mill A was found to have the highest precision for all features except for the 500µm channels and 500µm pillars for which mills D and C had the highest precision respectively (Mill A: 7.64 (500µm channel), 7.44 (500µm pillars) Mill C 6.25 (500µm channel) Mill D 4.61 (500µm pillars). For all the features tested, the 3D printer had a higher CV than the mills indicating that for the tested features a higher tolerance will be achieved using CNC machining. The 3D printer was found to have max CV of 23.6 and a minimum of 4.99 for the 200µm phase guide and 1000µm pillars, respectively. The highest CV found for any mill and any feature was 16.58 with mill B for the 200µm phase guide. They hobbyist unit performed well achieving a CVs down to 2.03 (1000µm channel) with a maximum of 12.12 (200µm phase guide). The specific application and acceptable tolerances should guide machine selection and manufacturing technique. These results support that the tested mills can create both simple and complex microfluidic devices directly into cell culture plates.

**Table 1.**
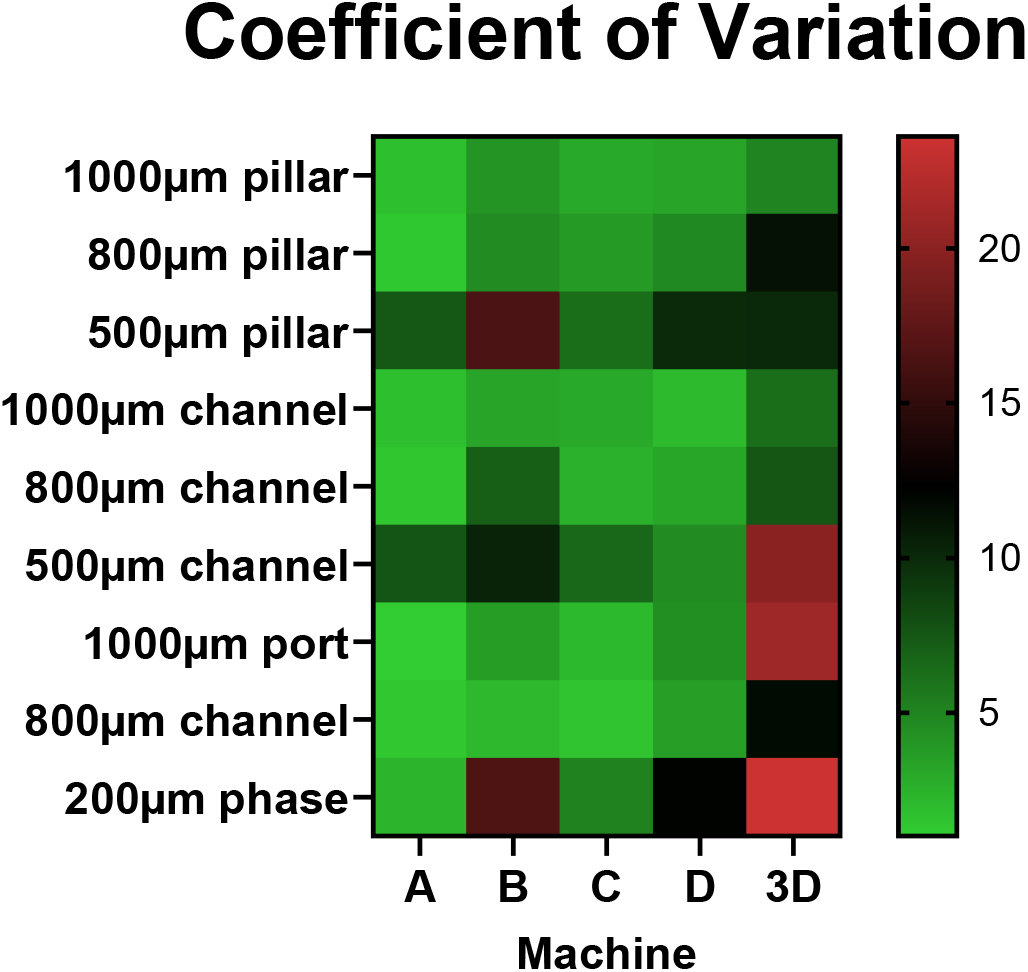
Coefficient of variation results for all machines. Heat map is plotted as a 3-color scale using the minimum CV (green), 50^th^ percentile CV (black), and maximum CV (red).

The variability between runs was assessed using the two independent trials conducted on mills B, C, and D. Differences were found for mills B, C, and D for the 9 features measured (6/9, 1/9, 3/9 respectively). The specific findings can be found in **Supplemental Table 1**. The large variability between trials for mill B is concerning and warrants discussion. This variation is potentially due to the condition of the machine and its’ maintenance history or lack thereof. This machine was purchased used from an industrial setting where it had been neglected. The variability between trials is especially evident in the 800µm and 500µm channels where two distinct populations can be seen (**Figure 4BV-VI**). The distinct populations seen also contribute to the high CV of these features (7.03 and 10.35 respectively).

The results of both milling tests demonstrate that like any manufacturing technique there are sources of variation and error that result in manufactured components that deviate slightly from nominal. The amount of acceptable deviation or tolerance is dependent on the application at hand. There are several potential differences that could have contributed to the variations seen in our testing. The tooling has a tolerance associated with it that can contribute to variation in the milling. One of the major differences is the components that make up the mills evaluated in this testing. Mill D and is a lightweight desktop hobby while mills A, B, and C are free standing mills that are comprised of heavy cast iron structures intended to minimize vibrations. While mill C and B are both manufactured by Syil and share the X5 name, they are quite different machines. They have completely different spindle units with mill C being fitted with a high-speed water-cooled spindle while mill B has a low speed (5000 RPM max) air cooled spindle. Another difference is the type of motor used to control the movement of the axes. Mill B is controlled with servo motors while mills C and D use stepper motors. Another potential source of variation is the time that each mill took to cut the test patterns. While each mill was run with the same code at the same feed rate, there are differences in the run time between all three mills suggesting differences in acceleration and/or feed rate. It is suspected that some or all these potential sources of variation cause the variability seen in our testing.

The dimensions of the features that can be cut into the bottom of a plate are limited by several factors including the thickness of the bottom of the plate, the radius (R) of the endmill (i.e. cutting tool), and the spatial accuracy of the CNC machine. Plates with ∼1.0-1.3mm of plastic on the bottom are largely suitable for bottom milling while imaging bottom and glass bottom plates are not. Sharp outer corners (90°) are achievable while inner corners are limited by the radius of the tool used to cut them. It is also recommended that inner corners be designed to give extra clearance (∼150% R) to reduce tool vibration (chatter) and promote a clean corner and finish. Similarly, ports should be designed with clearance (∼250% R) on the diameter to allow the endmill to spiral down as endmills are not well suited for vertical plunging. Use of a drill bit should be considered when a design incorporates multiple holes of the same diameter or holes of a size that is impractical to cut using a spiral approach. Channels can be cut at a width of 2R (diameter of endmill) with a depth that is limited by the thickness of the plate and the tolerance of the mill being used. Pillar arrays are another common microfluidic feature that can be incorporated via bottom milling. Pillar arrays with spacing equal to the diameter of the tool + 2R are achievable. Phase barriers are the final feature of focus for bottom milling. These features are useful for promoting laminar flow, separating phases, and promoting pinning of gels or droplets. There are no design constraints from a tool standpoint if the phase barrier is designed such that the tool can cut along both sides.

One of the main challenges with employing CNC machining to microplates is quality of the starting stock. The flatness of the stock, in this case the well plate, partially dictates what can be created. To demonstrate this, variation in flatness for the lot used in testing was conducted. Flatness was assessed both across the plate and across a single well. These findings are plotted in **Figure 5** as the normalized position in Z either across the plate (**5A**) or across well (**5B**). An average difference of 124µm (standard deviation 20.4µm) across the plates and an average difference of 26.0µm (standard deviation 3.6µm) across a single well was found. The variation across the well is likely a result of the manufacturing process of the plates where they are cooled too fast allowing heat sinks to form. These heat sinks are visualized by **Figure 5B** where the points at the edges the well are higher than those in the middle and an approximated curve of the surface of the well can be visualized. The data for the flatness across the plate show that there are variations in flatness between locations across a plate. These results illustrate the potential challenge of using this technique to create features of accurate depth. Those interested in holding tight tolerances in z should consider testing multiple brands and lots of plates to determine a stock source compatible with their requirements.

**Figure 5:**
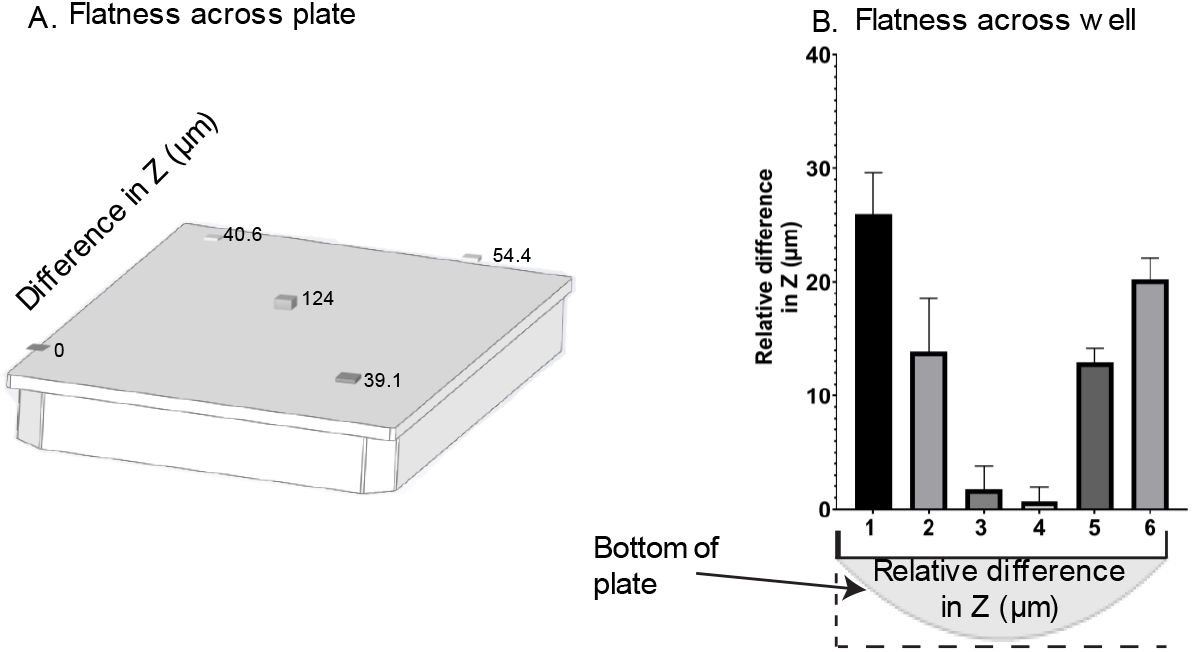
**A)** Plot of flatness across the plate. Relative Z values are plotted for the four corner wells and center well measured. **B)** Plot of flatness across a single well

Labs are already adopting this technology to create devices to facilitate the study of various physiologies. In **Table 2**, examples of labs already using this technology to create cellular MPMs are shown ^8-10, 13^. Microplate micromilling has been applied to study physiologies ranging from developmental toxicity to immune system modulation during cancer treatment. This technology is broadly useful and can be applied to a myriad of different physiologies and use cases making it well suited for NAMs development. The ability to create these devices using low-mid grade CNC machines facilitates adoption in a lab environment where space and noise are concerns. Our testing of noise levels while machining microplates demonstrates that levels remain below both the Michigan Occupational Safety and Health Administration (MIOSHA) Permissible Exposure Limit (PEL) of 90dBA and the Action Level of 85dBA (**Supplemental File 3**). While the applications to date have used micromilling to create cellular models the opportunity to expand both to other physiologies as well as to other research organisms is vast. Micromilling can facilitate the development of non-cellular NAMs including organisms commonly used in screening including *C. elegans, daphnia magna*, and zebra fish.

**Table 2.**
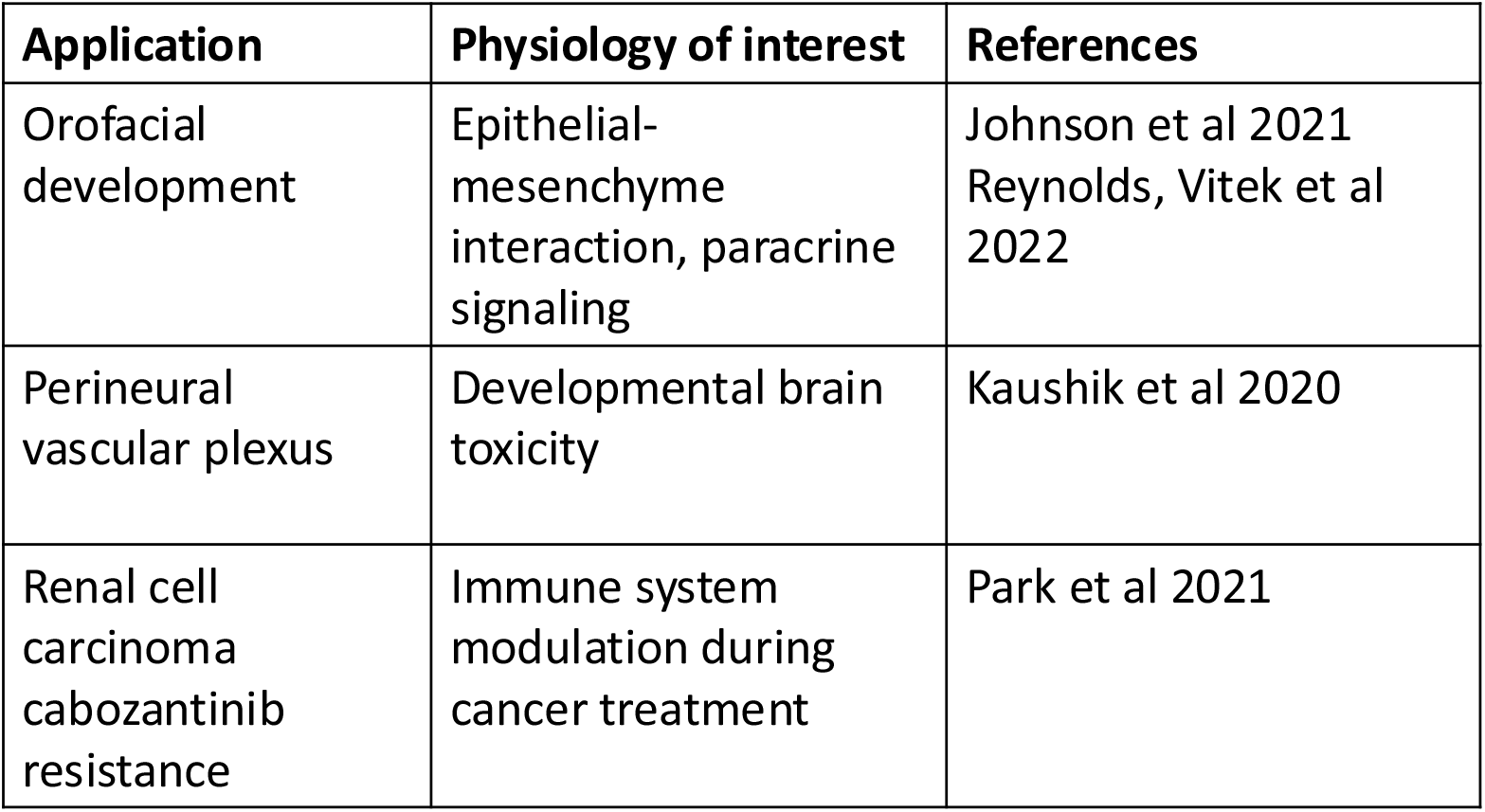
Example applications of micromilling used to create microphysiological systems.

| Application | Physiology of interest | References |
| --- | --- | --- |
| Orofacial development | Epithelial-mesenchyme interaction, paracrine signaling | Johnson et al 2021<br>Reynolds, Vitek et al 2022 |
| Perineural vascular plexus | Developmental brain toxicity | Kaushik et al 2020 |
| Renal cell carcinoma cabozantinib resistance | Immune system modulation during cancer treatment | Park et al 2021 |

## Conclusions

As we demonstrate with this work, CNC micromilling is a robust technology that can be applied to create microfluidic features directly into standard well plates and can be accomplished using a hobbyist grade machine. The ability to use an inexpensive and low-profile machine greatly lowers the bar of entry to life scientists seeking to adopt this technology in their research. Micromilling allows for the incorporation of microfluidic channels and features directly into standard well plates. These channels and features can be cut either into the top and/or the bottom of the plate, allowing for the creation of new microfluidic devices as well the re-creation of existing microfluidic devices into the well plate format. Milling into the bottom of the plate allows for more traditional microfluidic device designs to be created that are well suited for creating microphysiological models. We successfully demonstrate the ability of different CNC machines to create micro scale features including channels, pillars, and phase-guides. Differences in accuracy and precision were found between the tested mills and between milling and 3D printing with milling holding higher accuracy and precision. For the CNC mills, accuracy and precision correlated with the grade of the mill with the low-end hobbyist unit still performing at a level that would allow for successful MPM creation.

## Author Contributions

**Jacob Reynolds:** Conceptualization, Methodology, Investigation, Formal analysis, Writing-Original Draft, Writing-Review & Editing, Visualization **Meredith Adams:** Conceptualization, Methodology, Writing-Original Draft, Visualization **Jose Jimenez Brian Johnson:** Conceptualization, Methodology, Writing-Original Draft, Writing-Review & Editing, Visualization

## Acknowledgements

We thank Dhruv Singh for his assistance with data collection and review of the manuscript. We thank Robert Bennet and the Michigan State Department of Physics and Astronomy Machine Shop for their machining expertise and use of their Hurco mill. We thank Stephen Branch and the 3D Printing Core of the Institute of Quantitative Health Science and Engineering at Michigan State University for their 3D printing expertise and use of their 3D printer.

This work was supported by the National Institutes of Health R00-ES028744 to BPJ, National Institute of Environmental Health Sciences P42ES004911 to BPJ, and MSU Center for Research on Ingredient Safety (CRIS) Predoctoral fellowship to JIR.

## Notes and references

The data supporting this article have been included as part of the Supplementary Information.

Mill A is housed and maintained through the Physics and Astronomy Machine shop at Michigan State University. Mills B and C were both purchased used with mill C appearing to have been better maintained. Mill D was purchased new. Mills B,C, and D belong to the Johnson lab and are housed in the Institute for Quantitative Health Science and Engineering (IQ) at Michigan State University.

## Supplements

**Supplemental Figure 1:**
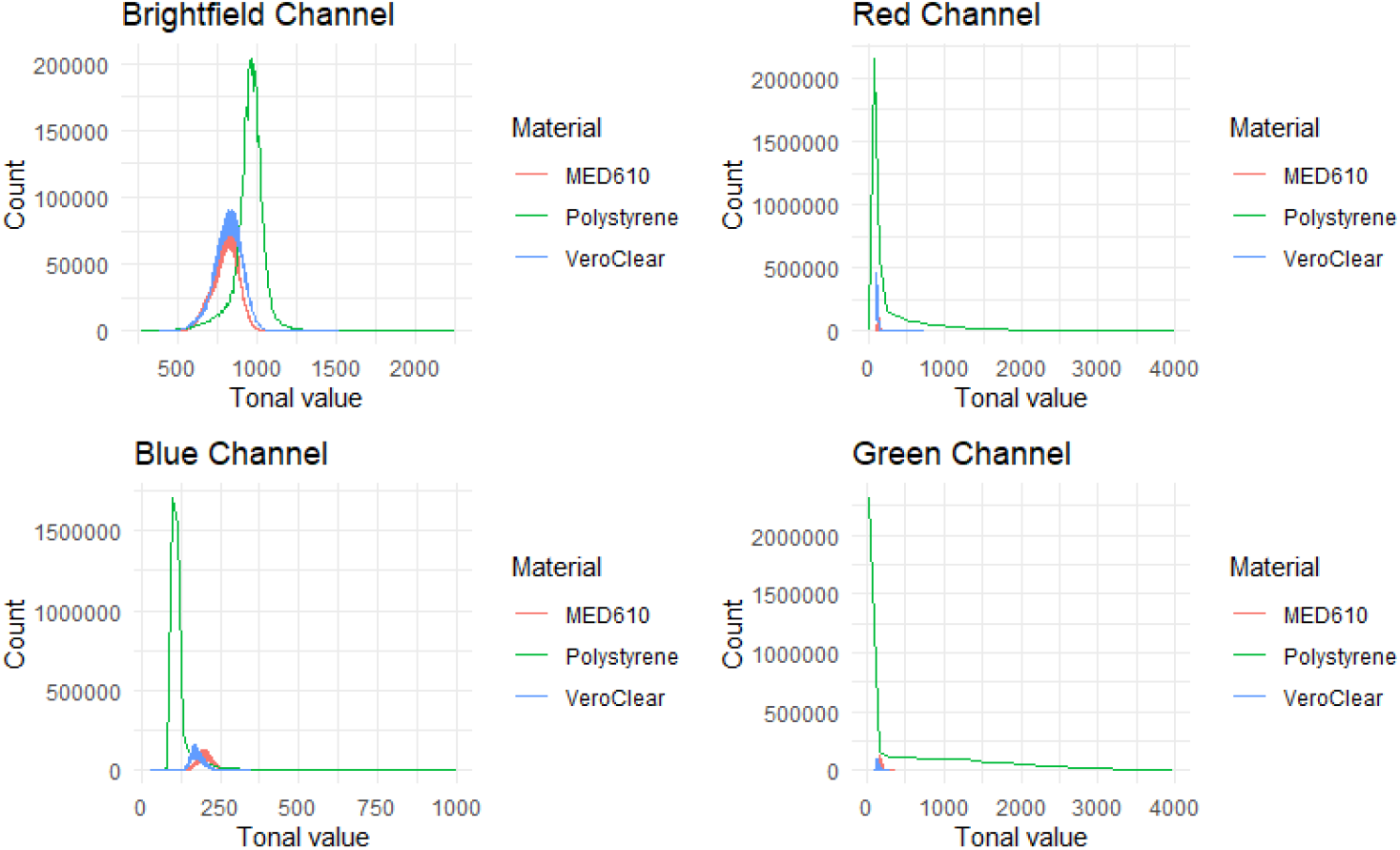
Image histograms for microscopy images for PS, VeroClear, and MED610 for the four channels imaged. **A) Brightfield B) Red C) Blue D) Green**

**Supplemental Table 1:** Comparison of trial 1 vs. trial 2 for mills B, C, D. 2way ANOVA followed by Sidak’s multiple comparisons test was performed using GraphPad Prism. Significance is reported as *P≤0.05, **P≤0.01, ***P≤0.001, ****P≤0.0001.

| Device | Feature | Machine |  |  |
| --- | --- | --- | --- | --- |
|  |  | B | C | D |
| Complex geometry | 1000µm pillar | ns | ns | ns |
|  | 800µm pillar | ns | ns | ns |
|  | 500µm pillar | *** | ns | ns |
|  | 1000µm channel | ** | ns | ns |
|  | 800µm channel | **** | ns | * |
|  | 500µm channel | **** | * | ns |
| Traditional | 1000µm port | ** | ns | * |
|  | 800µm channel | ns | ns | ** |
|  | 200µm phase | * | ns | ns |

**Supplemental Table 2:** All pairwise comparisons between machines. One-way ANOVA followed by Tukey’s multiple comparison test was performed using GraphPad Prism. Significance is reported as *P≤0.05, **P≤0.01, ***P≤0.001, ****P≤0.0001

| Device | Feature | Comparison |  |  |  |  |  |  |  |  |  |
| --- | --- | --- | --- | --- | --- | --- | --- | --- | --- | --- | --- |
|  |  | Expected vs. A |  | Expected vs. B |  | Expected vs.C |  | Expected vs. D |  | Expected vs. 3D |  |
|  |  | Summary | P value | Summary | P value | Summary | P value | Summary | P value | Summary | P value |
| Complex geometry | 1000µm pillar | ns | 0.9999 | **** | <0.0001 | *** | 0.0001 | ns | 0.1706 | ns | 0.0558 |
|  | 800µm pillar | ns | 0.9972 | *** | 0.0004 | ** | 0.0042 | ** | 0.0042 | ns | 0.8236 |
|  | 500µm pillar | ns | 0.361 | **** | <0.0001 | *** | 0.0001 | * | 0.0247 | **** | <0.0001 |
|  | 1000µm channel | ns | >0.9999 | ns | 0.243 | ns | 0.8429 | ** | 0.0037 | ns | 0.9993 |
|  | 800µm channel | ns | 0.9969 | ns | 0.3373 | ns | 0.6682 | * | 0.0305 | ns | 0.4512 |
|  | 500µm channel | ns | 0.1173 | **** | <0.0001 | ** | 0.0069 | **** | <0.0001 | **** | <0.0001 |
| Traditional | 1000µm port | ns | >0.9999 | ns | 0.2748 | ns | 0.5389 | ns | 0.9256 | ns | 0.6341 |
|  | 800µm channel | ns | 0.9998 | ns | 0.1231 | ** | 0.0046 | **** | <0.0001 | ** | 0.0028 |
|  | 200µm phase | ns | >0.9999 | ns | >0.9999 | ns | 0.4191 | ns | 0.2714 | **** | <0.0001 |
| Device | Feature | Comparison |  |  |  |  |  |  |  |  |  |
|  |  | A vs. B |  | A vs. C |  | A vs. D |  | A vs. 3D |  | B vs. C |  |
|  |  | Summary | P value | Summary | P value | Summary | P value | Summary | P value | Summary | P value |
| Complex geometry | 1000µm pillar | **** | <0.0001 | *** | 0.0003 | ns | 0.2725 | * | 0.029 | ns | 0.9841 |
|  | 800µm pillar | **** | <0.0001 | *** | 0.0008 | *** | 0.0008 | ns | 0.9738 | ns | 0.9835 |
|  | 500µm pillar | **** | <0.0001 | ns | 0.0852 | ns | 0.8454 | **** | <0.0001 | ** | 0.0079 |
|  | 1000µm channel | **** | <0.0001 | ** | 0.0062 | **** | <0.0001 | ns | 0.886 | * | 0.0252 |
|  | 800µm channel | *** | 0.0001 | * | 0.027 | **** | <0.0001 | **** | <0.0001 | ns | 0.5801 |
|  | 500µm channel | *** | 0.0001 | ns | 0.9116 | *** | 0.0009 | **** | <0.0001 | ** | 0.0044 |
| Traditional | 1000µm port | ns | 0.2847 | ns | 0.5519 | ns | 0.9315 | ns | 0.6469 | ns | 0.9977 |
|  | 800µm channel | ns | 0.2216 | * | 0.0113 | **** | <0.0001 | ** | 0.001 | ns | 0.8532 |
|  | 200µm phase | ns | 0.9991 | ns | 0.2969 | ns | 0.179 | **** | <0.0001 | ns | 0.5139 |
| Device | Feature | Comparison |  |  |  |  |  |  |  |  |  |
|  |  | B vs. D |  | B vs. 3D |  | C vs. D |  | C vs. 3D |  | D vs. 3D |  |
|  |  | Summary | P value | Summary | P value | Summary | P value | Summary | P value | Summary | P value |
| Complex geometry | 1000µm pillar | * | 0.0411 | **** | <0.0001 | ns | 0.2034 | **** | <0.0001 | **** | <0.0001 |
|  | 800µm pillar | ns | 0.9833 | **** | <0.0001 | ns | >0.9999 | **** | <0.0001 | **** | <0.0001 |
|  | 500µm pillar | **** | <0.0001 | **** | <0.0001 | ns | 0.6558 | **** | <0.0001 | **** | <0.0001 |
|  | 1000µm channel | **** | <0.0001 | **** | <0.0001 | **** | <0.0001 | *** | 0.0001 | **** | <0.0001 |
|  | 800µm channel | * | 0.0166 | **** | <0.0001 | **** | <0.0001 | **** | <0.0001 | **** | <0.0001 |
|  | 500µm channel | ns | 0.9923 | **** | <0.0001 | * | 0.0264 | **** | <0.0001 | **** | <0.0001 |
| Traditional | 1000µm port | ns | 0.8507 | ns | 0.9913 | ns | 0.9789 | ns | >0.9999 | ns | 0.9924 |
|  | 800µm channel | ns | 0.0573 | **** | <0.0001 | ns | 0.542 | **** | <0.0001 | **** | <0.0001 |
|  | 200µm phase | ns | 0.3502 | **** | <0.0001 | ns | 0.9998 | **** | <0.0001 | **** | <0.0001 |

**Supplemental file 1:** Machine code

**Supplemental file 2:** All measurement data used in support of this work

**Supplemental file 3:** Noise level recordings

