## Supplemental File 2 for "Microplate micromilling: A customizable platform to support the prototyping, development and testing of microphysiological culture models"

Supplemental file 2- All measurement data

Mill A-Hurco

| | Plate | | | | | | | | | | | | | | | | | | | |
|---|---|---|---|---|---|---|---|---|---|---|---|---|---|---|---|---|---|---|---|---|
| | 1 | | | 2 | | | 3 | | | 4 | | | 5 | | | 6 | | | Average | Standard deviation |
| diameter 1000um pillar | 991.25 | 981.29 | 999.448 | 980.11 | 992.948 | 1001.132 | 983.054 | 1006.241 | 981.36 | 983.596 | 1008.26 | 972.098 | 1023.311 | 1002.46 | 984.364 | 1001.744 | 1030.573 | 1030.332 | 997.4206 | 16.98843 |
| Diamteter 800um pillar 1 | 801.117 | 793.593 | 812.168 | 812.558 | 804.662 | 803.592 | 828.132 | 816.067 | 825.807 | 797.999 | 812.5 | 799.923 | 799.751 | 802.809 | 808.982 | 797.946 | 816.552 | 790.666 | 806.9347 | 10.12929 |
| Diameter 500um pillar 1 | 499.116 | 484.116 | 475.456 | 466.667 | 482.546 | 488.712 | 512.574 | 495.58 | 475.569 | 446.423 | 511.583 | 499.116 | 411.52 | 374.555 | 439.94 | 478.203 | 479.35 | 460.033 | 471.1699 | 34.06591 |
| width 1000um channel | 1027 | 1020.5 | 1027 | 1010.75 | 1010.75 | 1007.5 | 1017.25 | 1007.5 | 971.75 | 981.5 | 1001 | 988 | 1001 | 1001 | 984.75 | 975 | 994.5 | 978.25 | 1000.278 | 16.94146 |
| width 800um channel | 827.634 | 827.634 | 831.295 | 820.321 | 812.541 | 813.15 | 827.94 | 821.35 | 821.145 | 801.407 | 789.717 | 815.41 | 814.195 | 828.75 | 818.394 | 832.228 | 832.228 | 823.181 | 819.9178 | 10.84918 |
| width 500um channel | 520.822 | 491.825 | 497.292 | 543.916 | 555.218 | 545.003 | 504.012 | 507.843 | 510.757 | 591.83 | 594.386 | 597.311 | 574.965 | 591.152 | 587.307 | 501.554 | 495.068 | 491.277 | 538.9743 | 40.01851 |
| 1000um port | 1007.862 | 998.963 | 1024.781 | 1003.113 | 1002.023 | 1007.615 | 1007.699 | 996.102 | 1001.981 | 996.272 | 982.178 | 996.5 | 1004.587 | 983.741 | 985.243 | 1008.742 | 999.146 | 1004.587 | 1000.619 | 9.903394 |
| 800um channel | 793 | 793.007 | 786.5 | 806 | 809.25 | 786.5 | 809.25 | 799.5 | 786.5 | 806 | 806 | 819 | 815.75 | 819 | 812.5 | 806 | 806 | 806 | 803.6532 | 10.32289 |
| 200um phase | 198.25 | 191.75 | 195 | 204.75 | 198.25 | 204.75 | 208 | 208 | 208 | 208 | 204.75 | 208 | 201.5 | 201.5 | 201.5 | 201.5 | 201.5 | 204.75 | 202.7639 | 4.613892 |

Mill B-Syil Combo

| | | Plate | | | | | | | | | | |
|---|---|---|---|---|---|---|---|---|---|---|---|---|
| | Trial | 1 | | | 2 | | | 3 | | | Average | Standard Deviation |
| diameter | 1 | 876.904 | 972.315 | 937.95 | 896.111 | 1005.012 | 989.25 | 994.901 | 985.688 | 975.72 | 942.5984 | 37.24903 |
| 1000um | 2 | 933.209 | 931.662 | 943.967 | 950.251 | 897.33 | 936.406 | 913.967 | 891.709 | 934.419 | | |
| Diamteter | 1 | 705.033 | 702.579 | 733.125 | 711.401 | 764.911 | 729.659 | 756.077 | 793.593 | 790.365 | 734.4546 | 33.07838 |
| 800um | 2 | 744.278 | 703.721 | 742.211 | 769.749 | 763.093 | 734.133 | 704.531 | 700.923 | 670.8 | | |
| Diameter | 1 | 455.603 | 308.801 | 350.398 | 406.99 | 380.028 | 364.956 | 494.524 | 480.352 | 474.834 | 380.4566 | 60.64441 |
| 500um | 2 | 411.417 | 345.541 | 361.116 | 386.805 | 351.752 | 328.395 | 341.25 | 333.755 | 271.701 | | |
| width | 1 | 1075.77 | 1082.645 | 1059.998 | 1059.58 | 1069.566 | 1043.331 | 1030.25 | 1040.02 | 1036.832 | 1083.958 | 34.38898 |
| 1000um | 2 | 1108.25 | 1111.5 | 1111.5 | 1088.75 | 1085.5 | 1092 | 1137.5 | 1140.75 | 1137.5 | | |
| width | 1 | 983.698 | 971.163 | 976.953 | 959.411 | 945.208 | 957.493 | 868.96 | 856.824 | 894.341 | 796.5144 | 60.44392 |
| 800um | 2 | 856.38 | 851.302 | 852.083 | 816.442 | 819.78 | 829.814 | 838.324 | 818.929 | 830.183 | | |
| width | 1 | 689.008 | 682.531 | 676.008 | 667.042 | 660.038 | 653.541 | 604.579 | 624.008 | 633.758 | 611.0128 | 61.47814 |
| 500um | 2 | 637.414 | 645.802 | 647.109 | 518.22 | 508.757 | 517.771 | 553.531 | 534.896 | 544.217 | | |
| 1000um | 1 | 1057.909 | 1008.679 | 1037.499 | 1013.052 | 1058.228 | 1031.208 | 1043.498 | 1070.151 | 1023.151 | 1066.755 | 36.8766 |
| port | 2 | 1076.14 | 1145.1 | 1134.655 | 1100.5 | 1075.667 | 1091.78 | 1093.223 | 1077.143 | 1064 | | |
| 800um | 1 | 845 | 851.5 | 861.25 | 815.75 | 822.25 | 802.75 | 845 | 806 | 809.25 | 833.0833 | 17.33333 |
| channel | 2 | 841.75 | 854.75 | 858 | 832 | 835.25 | 822.25 | 825.5 | 832 | 835.25 | | |
| 200um | 1 | 126.75 | 143 | 172.25 | 178.75 | 172.25 | 172.25 | 195 | 191.75 | 191.75 | 178.425 | 31.90905 |
| phase | 2 | 237.75 | 230.5 | 227.5 | 230.75 | 230.75 | 237.25 | 221 | 204.75 | 201.5 | | |

Mill C- Syil Speedmaster

| | | Plate | | | | | | | | | | |
|---|---|---|---|---|---|---|---|---|---|---|---|---|
| | Trial | 1 | | | 2 | | | 3 | | | Average | Standard Deviation |
| diameter | 1 | 973.493 | 960.912 | 954.931 | 954.178 | 975.872 | 950.09 | 890.524 | 889.818 | 904.627 | 949.8436 | 27.2272 |
| 1000um | 2 | 950.34 | 969.448 | 939.52 | 966.6 | 987.749 | 958.684 | 942.59 | 959.521 | 968.287 | | |
| Diamteter | 1 | 755.574 | 735.535 | 726.896 | 779.953 | 790.445 | 758.651 | 721.302 | 711.817 | 682.856 | 744.5925 | 27.12669 |
| 800um | 2 | 721.566 | 728.464 | 728.384 | 774.837 | 748.672 | 784.409 | 754.427 | 745.101 | 753.776 | | |
| Diameter | 1 | 416.875 | 382.024 | 412.34 | 460.641 | 462.609 | 455.232 | 441.103 | 432.555 | 488.204 | 431.9328 | 26.21666 |
| 500um | 2 | 407.289 | 397.538 | 412.443 | 430.683 | 422.5 | 407.924 | 455.046 | 443.491 | 446.293 | | |
| width | 1 | 1033.5 | 1043.25 | 1030.25 | 981.522 | 1017.255 | 1001 | 1082.27 | 1069.27 | 1111.733 | 1044.962 | 29.96466 |
| 1000um | 2 | 1073.76 | 1062.75 | 1056.25 | 1072.5 | 1036.75 | 1040 | 1036.75 | 1033.5 | 1027 | | |
| width | 1 | 873.531 | 894.341 | 898.247 | 857.785 | 875.119 | 874.026 | 864.39 | 846.804 | 868.036 | 776.5456 | 21.42479 |
| 800um | 2 | 832.723 | 818.329 | 830.532 | 894.317 | 866.629 | 857.267 | 846.804 | 870.982 | 858.049 | | |
| width | 1 | 599.27 | 569.604 | 583.282 | 559.085 | 575.259 | 546.01 | 583.375 | 568.75 | 568.759 | 554.4521 | 35.44408 |
| 500um | 2 | 520.102 | 505.508 | 500.637 | 600.574 | 576.864 | 596.621 | 504.379 | 528.562 | 493.497 | | |
| 1000um | 1 | 1060.636 | 1083.63 | 1056.33 | 1025.93 | 1039.44 | 1022.47 | 1080.296 | 1078.001 | 1064.592 | 1052.811 | 21.47544 |
| port | 2 | 1052.649 | 1024.843 | 1027.185 | 1058.123 | 1078.648 | 1061.91 | 1034.935 | 1024.224 | 1076.751 | | |
| 800um | 1 | 838.5 | 845 | 825.5 | 858 | 858 | 861.25 | 851.5 | 861.25 | 877.5 | 848.25 | 11.91667 |
| channel | 2 | 838.5 | 854.75 | 851.5 | 841.75 | 841.75 | 848.25 | 841.75 | 835.25 | 838.5 | | |
| 200um | 1 | 165.75 | 172.25 | 175.5 | 191.86 | 198.25 | 188.528 | 175.53 | 165.75 | 175.53 | 158.4374 | 8.740838 |
| phase | 2 | 175.5 | 172.25 | 169 | 175.5 | 169 | 178.8 | 178.75 | 165.75 | 172.25 | | |

3D printer-Stratasys J750

| | Plate | | | | | | | | | | | | | | | | | | | |
|---|---|---|---|---|---|---|---|---|---|---|---|---|---|---|---|---|---|---|---|---|
| | 1 | | | 2 | | | 3 | | | 4 | | | 5 | | | 6 | | | Average | Standard deviation |
| diameter 1000um pillar | 960.165 | 953.907 | 900.76 | 1013.177 | 1075.647 | 1095.655 | 1048.502 | 1056.375 | 1021.483 | 1062.889 | 1044.666 | 1040 | 1006.162 | 1044.141 | 1033.367 | 1029.055 | 1050.293 | 1115.034 | 1030.627 | 49.98363 |
| Diamteter 800um pillar 1 | 895.928 | 880.318 | 817.845 | 807.185 | 818.329 | 796.091 | 626.846 | 657.657 | 618.705 | 882.673 | 865.337 | 900.96 | 814.577 | 850.197 | 828.399 | 926.204 | 833.908 | 906.395 | 818.1974 | 90.2952 |
| Diameter 500um pillar 1 | 572.757 | 659.197 | 574.662 | 561.799 | 615.796 | 589.031 | 581.786 | 702.068 | 565.173 | 651.591 | 590.356 | 590.973 | 665.012 | 643.344 | 636.287 | 533.683 | 462.278 | 508.009 | 594.6557 | 57.85606 |
| width 1000um channel | 1033.5 | 1046.5 | 1020.5 | 952.25 | 906.75 | 942.5 | 994.5 | 939.25 | 1056.25 | 942.5 | 1030.25 | 962 | 952.25 | 1131 | 913.25 | 1056.25 | 942.5 | 942.5 | 986.9167 | 59.72105 |
| width 800um channel | 763.197 | 728.595 | 690.998 | 738.351 | 735.858 | 707.463 | 667.208 | 744.165 | 671.776 | 606.088 | 670.493 | 760.812 | 667.232 | 739.316 | 728.5 | 768.884 | 792.047 | 837.189 | 723.2318 | 53.05704 |
| width 500um channel | 275.35 | 298.947 | 270.239 | 410.916 | 405.951 | 437.545 | 315.317 | 298.293 | 300.866 | 306.69 | 472.056 | 266.995 | 314.243 | 383.362 | 264.231 | 446.21 | 371.198 | 396.976 | 346.4103 | 66.84501 |
| 1000um port | 1351.695 | 1329.027 | 1314.303 | 969.579 | 1016.855 | 1003.298 | 1344.62 | 1302.111 | 1330.934 | 843.881 | 796.469 | 845.9 | 815.465 | 758.024 | 830.1 | 977.262 | 1002.961 | 1036.342 | 1048.268 | 213.7538 |
| 800um channel | 845 | 799.5 | 832 | 783.25 | 809.25 | 770.25 | 858 | 880.75 | 841.75 | 617.5 | 598 | 617.5 | 705.25 | 698.75 | 695.5 | 698.75 | 715 | 731.25 | 749.8472 | 85.12137 |
| 200um phase | 221.024 | 292.518 | 263.27 | 308.75 | 367.264 | 386.75 | 464.796 | 451.75 | 403.118 | 451.75 | 429.049 | 484.25 | 393.25 | 393.25 | 412.75 | 240.851 | 276.25 | 256.771 | 360.9673 | 82.78039 |

Coefficient of variation

| | | Machine | | | | |
|---|---|---|---|---|---|---|
| **Device** | **Feature** | **A** | **B** | **C** | **D** | **3D** |
| | **1000μm pi** | 1.75 | 4.07 | 2.95 | 3.21 | 4.99 |

| | | | | | | |
|---|---|---|---|---|---|---|
| | 800µm pill | 1.29 | 4.63 | 3.75 | 4.72 | 11.36 |
| | 500µm pill | 7.44 | 16.40 | 6.25 | 9.99 | 10.01 |
| | 1000µm cl | 1.74 | 3.26 | 2.95 | 2.03 | 6.23 |
| | 800µm ch | 1.36 | 7.03 | 2.56 | 3.15 | 7.55 |
| plex geon | 500µm ch | 7.64 | 10.35 | 6.58 | 4.61 | 19.86 |
| | 1000µm p | 1.02 | 3.56 | 2.10 | 4.42 | 20.98 |
| | 800µm ch | 1.32 | 2.14 | 1.45 | 3.50 | 11.68 |
| Traditiona | 200µm ph | 2.34 | 16.58 | 5.11 | 12.12 | 23.60 |

Plate flatness

| plate numb | a1 | a12 | h12 | h1 | e6 | e71 | e72 | e73 | e74 | e75 | e76 | difference | aver | std | across well aver | | std |
|---|---|---|---|---|---|---|---|---|---|---|---|---|---|---|---|---|---|
| 1 | 216736 | 216941 | 217086 | 216988 | 217786 | 218022 | 217905 | 217763 | 217723 | 217841 | 217912 | 105 | 124 | 20.4245 | 29.9 | 25.96667 | 3.611556 |
| 2 | 216227 | 216797 | 216925 | 216751 | 217683 | 217939 | 217832 | 217699 | 217687 | 217813 | 217881 | 145.6 | | | 25.2 | | |
| 3 | 216202 | 216644 | 216787 | 216598 | 217416 | 217668 | 217530 | 217440 | 217461 | 217583 | 217664 | 121.4 | | | 22.8 | | |

normal

| | | | | |
|---|---|---|---|---|
| 0 | 20.5 | 35 | 25.2 | 105 |
| 0 | 57 | 69.8 | 52.4 | 145.6 |
| 0 | 44.2 | 58.5 | 39.6 | 121.4 |

standard deviation

| | | | | |
|---|---|---|---|---|
| 0 | 18.51927 | 17.75284 | 13.60784 | 20.4245 |

normal

| | | | | | |
|---|---|---|---|---|---|
| 29.9 | 18.2 | 4 | 0 | 11.8 | 18.9 |
| 25.2 | 14.5 | 1.2 | 0 | 12.6 | 19.4 |
| 22.8 | 9 | 0 | 2.1 | 14.3 | 22.4 |

| | 1 | 2 | 3 | 4 | 5 | 6 | 7 | 8 | 9 | 10 | 11 | 12 |
|---|---|---|---|---|---|---|---|---|---|---|---|---|
| A | 0 | | | | | | | | | | | 40.6 |
| B | | | | | | | | | | | | |
| C | | | | | | | | | | | | |
| D | | | | | | | | | | | | |
| E | | | | | | 124 | | | | | | |
| F | | | | | | | | | | | | |
| G | | | | | | | | | | | | |
| H | 39.1 | | | | | | | | | | | 54.4 |

| Cell viability | | | | | | | | | |
|---|---|---|---|---|---|---|---|---|---|
| day | 1 | | | 2 | | | 3 | | |
| Plate | 1 | 2 | 3 | 1 | 2 | 3 | 1 | 2 | 3 |
| | 115472 | 153001.7 | 150963 | 225558.3 | 216285.7 | 194365.3 | 333003.3 | 329457 | 299389.7 |
| | 139574 | 159457.7 | 195889 | 135274.3 | 226269.7 | 221912.3 | 324353.3 | 320531 | 292192.7 |
| | 126412 | 145176.7 | 185502 | 174021.3 | 222118.7 | 231835.3 | 304519.3 | 317194 | 306517.7 |
| | 141383 | 142315.7 | 186590 | 205999.3 | 239599.7 | 249965.3 | 310364.3 | 316472 | 294001.7 |
| | 149828 | 139861.7 | 186949 | 135377.3 | 211128.7 | 224753.3 | 274754.3 | 200987 | 259637.7 |
| | 137866 | 137137.7 | 178338 | 233731.3 | 227523.7 | 242765.3 | 258162.3 | 305768 | 270808.7 |
| avg | 153984.2778 | | | 212138.0556 | | | 295450.7778 | | |
| std | 23157.09627 | | | 33053.47104 | | | 32665.36673 | | |
| n | 18 | | | 18 | | | 18 | | |

| Veroclear | | | | | | | | | |
|---|---|---|---|---|---|---|---|---|---|
| day | 1 | | | 2 | | | 3 | | |
| Plate | 1 | 2 | 3 | 1 | 2 | 3 | 1 | 2 | 3 |
| | 208920.7 | 300307.3 | 82851.67 | 306011.3 | 287025 | 49719.67 | 272325 | 238582.7 | 116889.7 |
| | 297700.7 | 308258.3 | 85093.67 | 315057.3 | 290341 | 65832.67 | 286073 | 255548.7 | 102872.7 |
| | 270511.7 | 280372.3 | 91029.67 | 322633.3 | 284678 | 67962.67 | 250095 | 252281.7 | 102102.7 |
| | 228736.7 | 292563.3 | 81570.67 | 311688.3 | 259090 | 75823.67 | 255886 | 245913.7 | 145877.7 |
| | 243111.7 | 341110.3 | 85788.67 | 332354.3 | 347740 | 72504.67 | 278094 | 258712.7 | 143712.7 |
| | 244583.7 | 318518.3 | 77049.67 | 294605.3 | 276352 | 64103.67 | 267647 | 188427.7 | 12764.67 |
| scale for surface area | | | | | | | | | |
| | 108012 | 155258.9 | 42834.31 | 158207.9 | 148391.9 | 25705.07 | 140792 | 123347.2 | 60431.96 |
| | 153911.2 | 159369.6 | 43993.43 | 162884.6 | 150106.3 | 34035.49 | 147899.7 | 132118.7 | 53185.17 |
| | 139854.5 | 144952.5 | 47062.34 | 166801.4 | 147178.5 | 35136.7 | 129299.1 | 130429.6 | 52787.08 |
| | 118256.9 | 151255.2 | 42172.03 | 161142.9 | 133949.5 | 39200.84 | 132293.1 | 127137.4 | 75418.75 |
| | 125688.7 | 176354 | 44352.74 | 171827.2 | 179781.6 | 37484.91 | 143774.6 | 133754.4 | 74299.45 |
| | 126449.8 | 164674 | 39834.68 | 152311 | 142874 | 33141.6 | 138373.5 | 97417.1 | 6599.333 |
| avg | 110238.1579 | | | 115564.5217 | | | 105519.9 | | |
| std | 51376.02928 | | | 60227.81406 | | | 41344.92 | | |
| n | 18 | | | 18 | | | 18 | | |

| MED610 | | | | | | | | | |
|---|---|---|---|---|---|---|---|---|---|
| day | 1 | | | 2 | | | 3 | | |
| Plate | 1 | 2 | 3 | 1 | 2 | 3 | 1 | 2 | 3 |
| | 288842 | 304315 | 302445.7 | 216285.7 | 263760.7 | 301984.7 | 278915.7 | 167757 | 268026.7 |
| | 285450 | 311457 | 300014.7 | 226269.7 | 274341.7 | 276578.7 | 237562.7 | 193408 | 267310.7 |
| | 288225 | 291739 | 329250.7 | 222118.7 | 287720.7 | 266330.7 | 235461.7 | 144618 | 242004.7 |
| | 276328 | 266907 | 299940.7 | 239599.7 | 294316.7 | 293323.7 | 228285.7 | 188794 | 263239.7 |
| | 281325 | 249416 | 284227.7 | 211128.7 | 266754.7 | 275086.7 | 246516.7 | 172348 | 210412.7 |
| | 309139 | 282180 | 282529.7 | 227523.7 | 292586.7 | 259522.7 | 237318.7 | 185607 | 280590.7 |
| scale for surface area | | | | | | | | | |
| | 149331.3 | 157330.9 | 156364.4 | 111819.7 | 136364.3 | 156126.1 | 144199.4 | 86730.37 | 138569.8 |
| | 147577.7 | 161023.3 | 155107.6 | 116981.4 | 141834.6 | 142991.2 | 122819.9 | 99991.94 | 138199.6 |
| | 149012.3 | 150829.1 | 170222.6 | 114835.4 | 148751.6 | 137693 | 121733.7 | 74767.51 | 125116.4 |
| | 142861.6 | 137990.9 | 155069.3 | 123873 | 152161.7 | 151648.3 | 118023.7 | 97606.5 | 136094.9 |
| | 145445 | 128948.1 | 146945.7 | 109153.5 | 137912.2 | 142219.8 | 127449.1 | 89103.92 | 108783.3 |
| | 159824.9 | 145887.1 | 146067.8 | 117629.7 | 151267.3 | 134173.2 | 122693.8 | 95958.82 | 145065.4 |
| avg | 150324.4136 | | | 134857.5543 | | | 116272.6681 | | |
| std | 9330.289683 | | | 15367.131 | | | 21304.10562 | | |
| n | 18 | | | 18 | | | 18 | | |
