## Supplementary figures and images for "Microplate micromilling: A customizable platform to support the prototyping, development and testing of microphysiological culture models"

### Supplemental File 3

Supplemental file 3: Noise level recordings

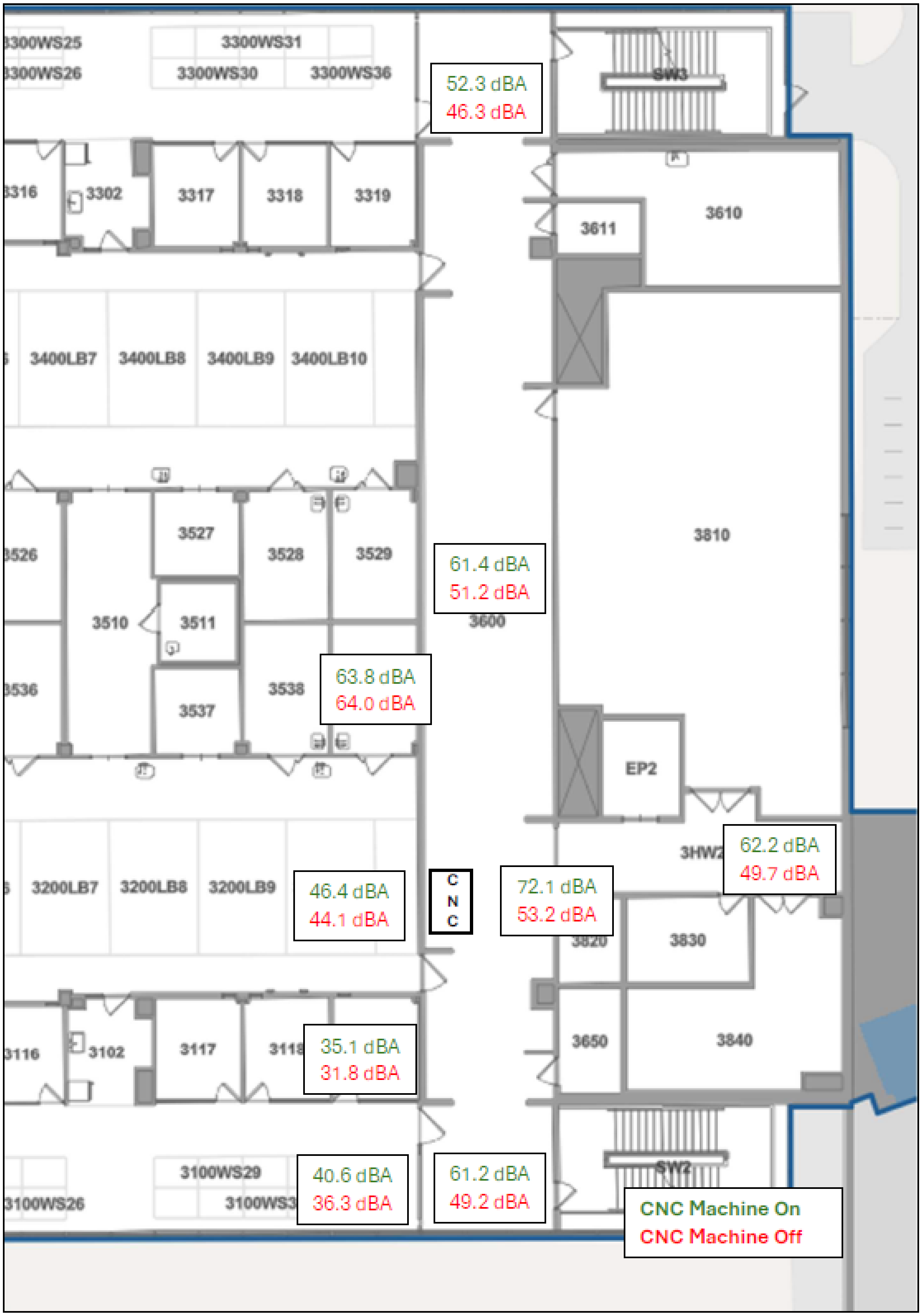
